# Multi-study inference of regulatory networks for more accurate models of gene regulation

**DOI:** 10.1101/279224

**Authors:** Dayanne M. Castro, Nicholas R. de Veaux, Emily R. Miraldi, Richard Bonneau

## Abstract

Gene regulatory networks are composed of sub-networks that are often shared across biological processes, cell-types, and organisms. Leveraging multiple sources of information, such as publicly available gene expression datasets, could therefore be helpful when learning a network of interest. Integrating data across different studies, however, raises numerous technical concerns. Hence, a common approach in network inference, and broadly in genomics research, is to separately learn models from each dataset and combine the results. Individual models, however, often suffer from under-sampling, poor generalization and limited network recovery. In this study, we explore previous integration strategies, such as batch-correction and model ensembles, and introduce a new multitask learning approach for joint network inference across several datasets. Our method initially estimates the activities of transcription factors, and subsequently, infers the relevant network topology. As regulatory interactions are context-dependent, we estimate model coefficients as a combination of both dataset-specific and conserved components. In addition, adaptive penalties may be used to favor models that include interactions derived from multiple sources of prior knowledge including orthogonal genomics experiments. We evaluate generalization and network recovery using examples from *Bacillus subtilis* and *Saccharomyces cerevisiae*, and show that sharing information across models improves network reconstruction. Finally, we demonstrate robustness to both false positives in the prior information and heterogeneity among datasets.

## Introduction

Gene regulatory network inference aims at computationally deriving and ranking regulatory hypotheses on transcription factor-target gene interactions [1–3]. Often, these regulatory models are learned from gene expression measurements across a large number of samples. Strategies to obtain such data range from combining several publicly available datasets to generating large expression datasets from scratch [4–7]. Given decreasing costs of sequencing and the exponential growth in the availability of gene expression data in public databases [8, 9], data integration across several studies becomes particularly promising for an increasing number of biological systems.

In theory, multi-study analyses provide a better representation of the underlying cellular regulatory network, possibly revealing insights that could not be uncovered from individual studies [6]. In practice, however, biological datasets are highly susceptible to batch effects [10], which are systematic sources of technical variation due to different reagents, machines, handlers etc. that complicate omics meta-analyses [11, 12]. Although several methods to remove batch effects from expression data have been developed, they often rely on evenly distributed experimental designs across batches [13, 14]. Batch-correction methods may deflate relevant biological variability or induce incorrect differences between experimental groups when conditions are unbalanced across batches, which can significantly affect downstream analyses [15]. Therefore these batch effect removal methods are not applicable when integrating public data from multiple sources with widely differing experimental designs.

In network inference, an approach often taken to bypass batch effects is to learn models from each dataset separately and combine the resulting networks [16, 17]. Known as ensemble learning, this idea of synthesizing several weaker models into a stronger aggregate model is commonly used in machine learning to prevent overfitting and build more generalizable prediction models [18]. In several scenarios, ensemble learning avoids introducing additional artifacts and complexity that may be introduced by explicitly modeling batch effects. On the other hand, the relative sample size of each dataset is smaller when using ensemble methods, likely decreasing the ability of an algorithm to detect relevant interactions. As regulatory networks are highly context-dependent [19], for example, TF binding to several promoters is condition-specific [20], a drawback for both batch-correction and ensemble methods is that they produce a single network model to explain the data across datasets. Relevant dataset-specific interactions might not be recovered, or just difficult to tell apart using a single model.

Although it will not be the primary focus of this paper, most modern network inference algorithms integrate multiple data-types to derive prior or constraints on network structure. These priors/constraints have been shown to dramatically improve network model selection performance when combined with the state variables provided by expression data. In these methods [17, 21], priors or constraints on network structure (derived from multiple sources like known interactions, ATAC-seq, DHS, or ChIP-seq experiments [22–24]) are used to influence the penalty on adding model components, where edges in the prior are effectively penalized less. Here we describe a method that builds on that work (and similar work in other fields), but in addition we let model inference processes (each carried out using a separate data-set) influence each others model penalties, so that edges that agree across inference tasks are more likely to be uncovered [25–31]. Several previous works on this front focused on enforcing similarity across models by penalizing differences on strength and direction of regulatory interactions using a fusion penalty [25, 27, 28]. Because the influence of regulators on the expression of targets may vary across datasets, possibly even due to differences in measurement technologies, we look to induce similarity on network structure (the choice of regulators) using a group-sparse penalty. Previous methods also applied this type of penalty [26, 29, 31], however, they were not robust to differences in relevant edges across datasets.

Here we propose a multitask learning (MTL) approach to exploit cross-dataset commonalities while recognizing differences and is able to incorporate prior knowledge on network structure if available [32, 33]. In this framework, information flow across datasets leads the algorithm to prefer solutions that better generalize across domains, thus reducing chances of overfitting and improving model predictive power [34]. Since biological datasets are often under-sampled, we hypothesize that sharing information across models inferred from multiple datasets using a explicit multitask learning framework will improve accuracy of inferred network models in a variety of common experimental designs/settings.

In this paper, we explicitly show that joint inference significantly improves network recovery using examples from two model organisms, *Bacillus subtilis* and *Saccharomyces cerevisiae*. We show that models inferred for each dataset using our MTL approach (which adaptively penalizes conserved and data-set-unique model components separately) are vastly more accurate than models inferred separately using a single-task learning (STL) approach. We also explore commonly used data integration strategies, and show that MTL outperforms both batch-correction and ensemble approaches. In addition, we also demonstrate that our method is robust to noise in the input prior information. Finally, we look at conserved and dataset-specific inferred interactions, and show that our method can leverage cross-dataset commonalities, while being robust to differences.

## Results

### Overview of network inference algorithm

To improve regulatory network inference from expression data, we developed a framework that leverages training signals across related expression datasets. For each gene, we assume that its regulators may overlap across conditions in related datasets, and thus we could increase our ability to uncover accurate regulatory interactions by inferring them jointly. Our method takes as input multiple expression datasets and priors on network structure, and then outputs regulatory hypotheses associated with a confidence score proportional to our belief that each prediction is true (Fig 1A). As previous studies [17, 35–37], our method also includes an intermediate step that estimates transcription factor activities (TFA), and then, models gene expression as a function of those estimates (Fig 1B).

**Fig 1:**
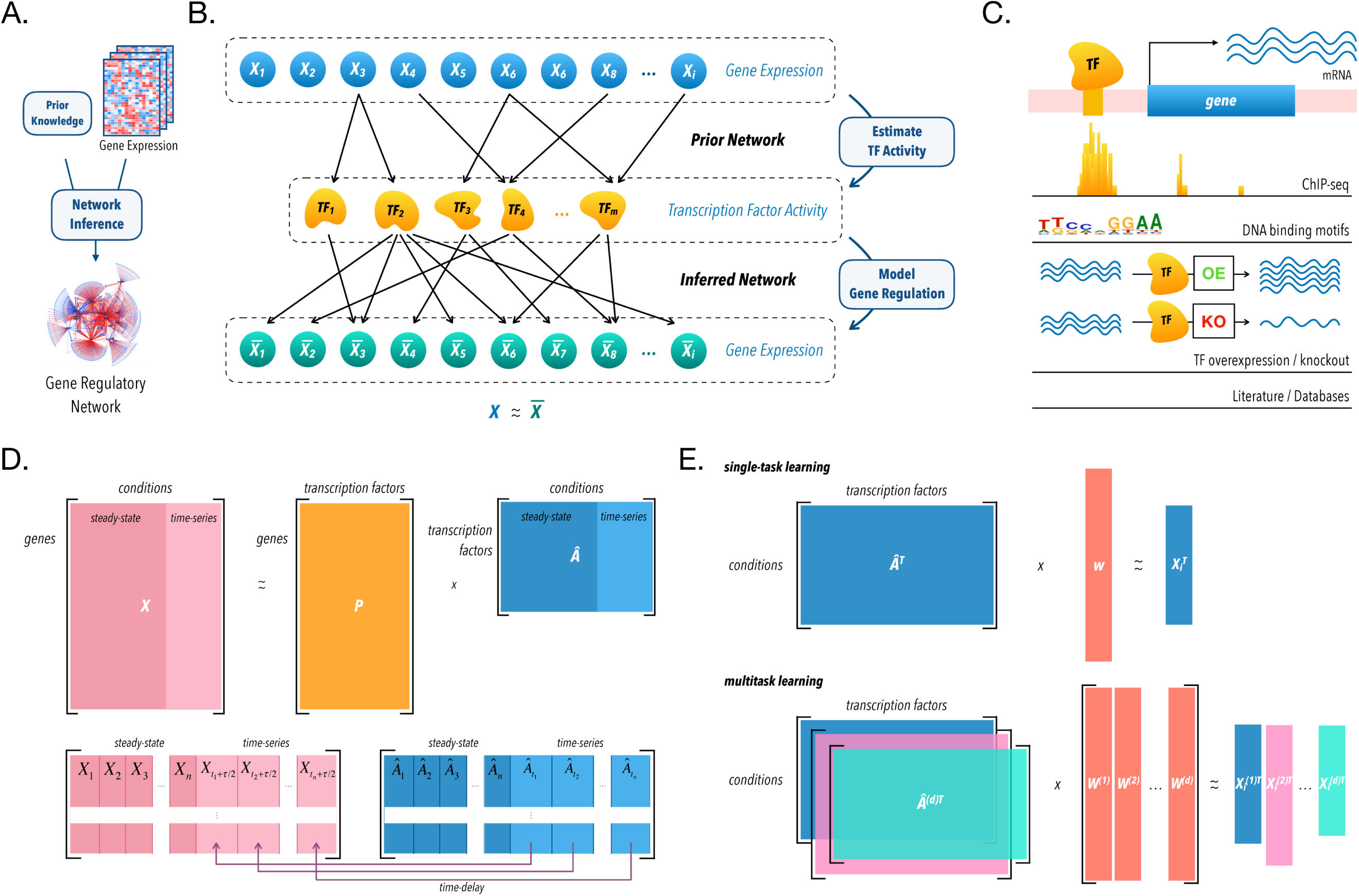
Gene regulatory network inference schematic. (A) Our network inference algorithm takes as input a gene expression matrix, *X*, and a prior on network structure and outputs regulatory hypotheses of regulator-target interactions. (B) Using priors on network topology and gene expression data, we estimate transcription factor activities (TFA), and subsequently model gene expression as a function of these activities. (C) We use several possible sources of prior information on network topology. (D) Prior information is encoded in a matrix *P*, where positive and negative entries represent known activation and repression respectively, whereas zeros represent absence of known regulatory interaction. To estimate hidden activities, we consider *X* = *PA* (top), where the only unknown is the activities. Of note, a time-delay is implemented for time-series experiments (bottom). (E) Finally, for each gene, we find regulators that influence its expression using regularized linear regression. We either learn these influences, or weights, for each dataset independently, single-task learning (top), or jointly through multi-task learning (bottom).

In our model, TFA represent a relative quantification of active protein that is inducing or repressing the transcription of its targets in a given sample, and is an attempt to abstract away unmeasured factors that influence TFA in a living cell [37–39], such as post-translational regulation [40], protein-protein interactions [41], and chromatin accessibility [42]. We estimate TFA from partial knowledge of the network topology (Fig 1C) [21, 43–47] and gene expression data as previously proposed (Fig 1D) [17]. This is comparable to using a TF’s targets collectively as a reporter for its activity.

Next, we learn the dependencies between gene expression and TFA and score predicted interactions. In this step, our method departs from previous work, and we employ multitask learning to learn regulatory models across datasets jointly, as opposed to single-task learning, where network inference is performed for each dataset independently (Fig 1E). As genes are known to be regulated by a small number of TFs [48], we can assume that these models are sparse, that is, they contain only a few nonzero entries [3]. We thus implement both approaches using sparsity-inducing penalties derived from the lasso [49]. Here the network model is represented as a matrix for each target gene (where columns are data-sets/cell-types/studies and rows are potential regulators) with signed entries corresponding to strength and type of regulation.

Importantly, our MTL approach decomposes this model coefficients matrix into a dataset-specific component and a conserved component to enable us to penalize dataset-unique and conserved interactions separately for each target gene [32]; this separation captures differences in regulatory networks across datasets (Fig 2). Specifically, we apply an *l*_1_*/l*_*∞*_ penalty to the one component to encourage similarity between network models [50], and an *l*_1_*/l*_1_ penalty to the other to accommodate differences [32]. We also incorporate prior knowledge by using adaptive weights when penalizing different coefficients in the *l*_1_*/l*_1_ penalty [33]. Finally, we perform this step for several bootstraps of the conditions in the expression and activities matrices, and calculate a confidence score for each predicted interaction that represents both the stability across bootstraps and the proportion of variance explained of the target expression dependent on each predictor.

**Fig 2:**
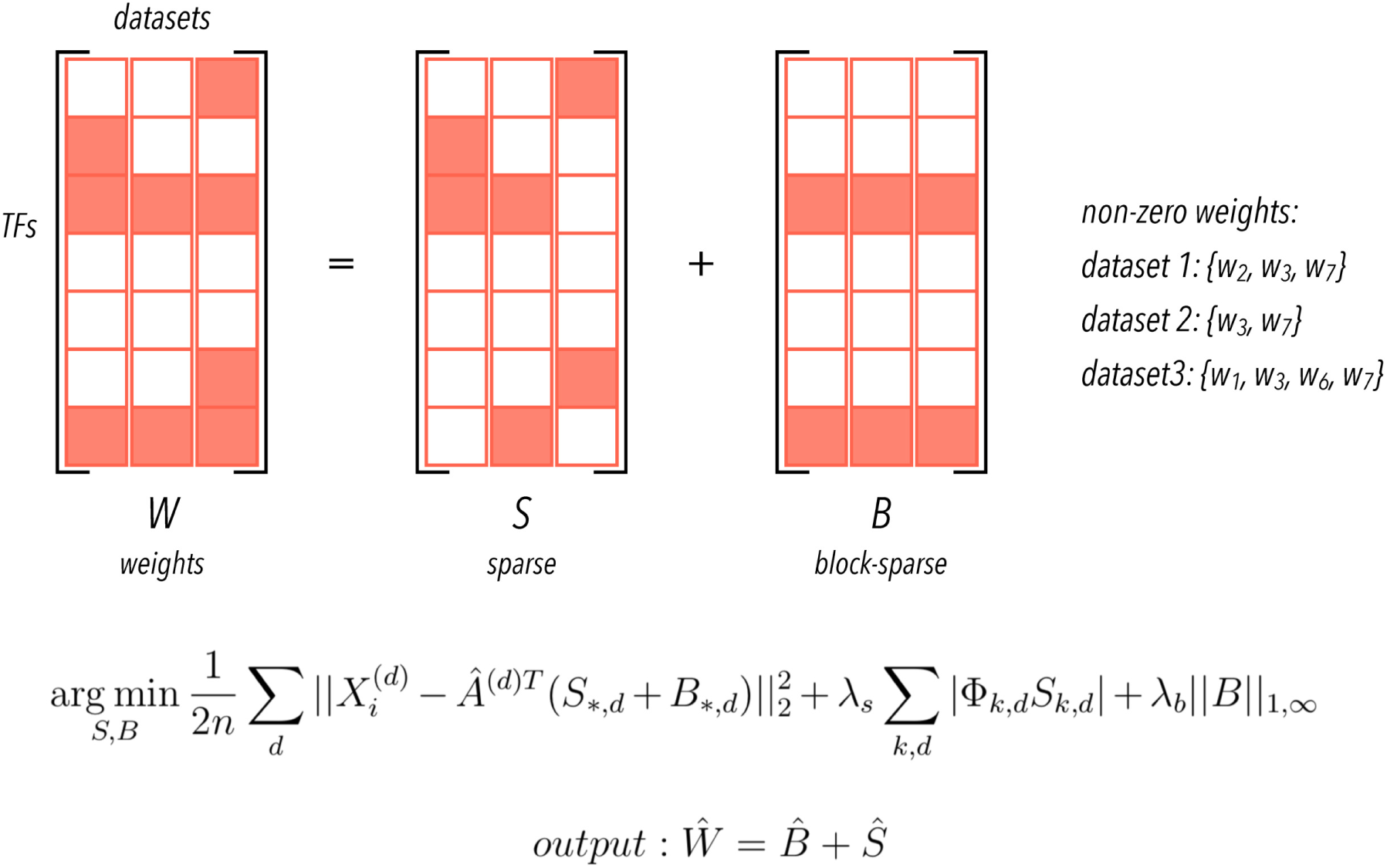
Representation of the weights matrix for one gene in the multitask setting. We represent model coefficients as a matrix *W* (predictors by datasets) where nonzero rows represent predictors relevant for all datasets. We decompose the weights into two components, and regularize them differently, using a sparse penalty (*l*_1_*/l*_1_ to *S* component) to encode a dataset-specific component and a block-sparse penalty (*l*_1_*/l*_*∞*_ to *B* component) to encode a conserved one. To illustrate, in this example, non-zero weights are shown on the right side. Note that, in this schematic example, regulators w3 and w7 are shared between all datasets. We also show the objective function minimized to estimate S and B on the bottom (for details, see methods).

Our method is readily available in an open-source package, *Inferelator-AMuSR* (**A**daptive **Mu**ltiple **S**parse **R**egression), enabling TF activity estimation and multi-source gene regulatory network inference, ultimately facilitating mechanistic interpretations of gene expression data to the Biology community. In addition, this method allows for adaptive penalties to favor interactions with prior knowledge proportional to the user-defined belief that interactions in the prior are true. Finally, our implementation also includes several mechanisms that speed-up computations, making it scalable for the datasets used here, and support for parallel computing across multiple nodes and cores in several computing environments.

### Model organisms, expression datasets, and priors

We validated our approach using two model organisms, a gram-positive bacteria, *B. subtilis*, and an eukaryote, *S. cerevisiae*. Availability of validated TF-target regulatory interactions, hereafter referred to as the gold-standard, make both organisms a good choice for exploring inference methods (3040 interactions, connecting 153 TFs to 1822 target genes for *B. subtilis* [17, 46], 1198 interactions connecting 91 TFs to 842 targets for S. cerevisiae [51]). For *B. subtilis*, we use two expression datasets. The first one, *B. subtilis 1*, was collected for strain PY79 and contains multiple knockouts, competence and sporulation-inducing conditions, and chemical treatments (429 samples, 38 experimental designs with multiple time-series experiments) [17]. The second dataset, *B. subtilis 2*, was collected for strain BSB1 and contains several nutritional, and other environmental stresses, as well as competence and sporulation-inducing conditions (269 samples, and 104 conditions) [52]. For *S. cerevisiae*, we downloaded three expression datasets from the SPELL database [53]. *S. cerevisiae 1* is a compendium of steady-state chemostat cultures with several combinations of cultivation parameters (170 samples, 55 conditions) [54]. *S. cerevisiae 2* profiles two yeast strains (BY and RM) grown with two carbon sources, glucose and ethanol, in different concentrations (246 samples, and 109 conditions) [55]. Finally, *S. cerevisiae 3* with expression profiles following several mutations and chemical treatments (300 samples) [56]. Each dataset was collected using a different microarray platform. Cross-platform data aggregation is well known to cause strong batch effects [10]. For each species, we considered the set of genes present across datasets.

In our inference framework, prior knowledge on network topology is essential to first estimate transcription factor activities and to then bias model selection towards interactions with prior information during the network inference stage of the algorithm. Therefore, to properly evaluate our method, it is necessary to gather prior interactions independent of the ones in the gold-standard. For *B. subtilis*, we adopt the previously used strategy of partitioning the initial gold-standard into two disjoint sets, a prior for use in network inference and a gold-standard to evaluate model quality [17]. For *S. cerevisiae*, on the other hand, we wanted to explore a more realistic scenario, where a gold-standard is often not available. In the absence of such information, we hypothesized that orthogonal high-throughput datasets would provide insight. Because the yeast gold-standard [51] was built as a combination of TF-binding (ChIP-seq, ChIP-ChIP) and TF knockout datasets available in the YEASTRACT [47] and the SGD [57] databases, we propose to derive prior knowledge from chromatin accessibility data [22, 23] and TF binding sites [58] (as this is a realistic and efficient genomic experimental design for non-model organisms). Open regions in the genome can be scanned for transcription factor binding sites, which can provide indirect evidence of regulatory function [59]. We then assigned TFs to the closest downstream gene, and built a prior matrix where entries represent the number of motifs for a particular TF that was associated to a gene [60, 61]. We obtained a list of regulators from the YeastMine database [62], which we also used to sign entries in the prior: interactions for regulators described as repressors were marked as negative. Because genome-wide measurements of DNA accessibility can be obtained in a single experiment, using techniques that take advantage of the sensitivity of nucleosome-free DNA to endonuclease digestion (DNase-seq) or to Tn5 transposase insertion (ATAC-seq) [63], we expect this approach to be generalizable to several biological systems.

### Sharing information across network models via multitask learning improves model accuracy

Using the above expression datasets and priors, we learn regulatory networks for each organism employing both single-task and our multitask approaches. To provide an intuition for cross-dataset transfer of knowledge, we compare confidence scores attributed to a single gold-standard interaction using either STL or MTL for each organism. For *B. subtilis*, we look at the interaction between the TF *sigB* and the gene *ydfP* (Fig 3A). The relationship between the *sigB* activity and *ydfP* expression in the first dataset *B. subtilis 1* is weaker than in *B. subtilis 2*. This is reflected in the predicted confidence scores, a quarter as strong for *B. subtilis 1* than for *B. subtilis 2*, when each dataset is used separately to learn networks through STL. On the other hand, when we learn these networks in the MTL framework, information flows from *B. subtilis 2* to *B. subtilis 1*, and we assign a high confidence score to this interaction in both networks. Similarly, for *S. cerevisiae*, we look at the interaction between the TF *Rap1* and the target gene *Rpl12a* (Fig 3B). In this particular case, we observe a strong and easier-to-uncover relationship between *Rap1* estimated activity and *Rpl12a* expression for all datasets. Indeed, we assign a nonzero confidence score to this interaction for all datasets using STL, although for *S. cerevisiae 2 and 3* these are much smaller than the scores attributed when networks are learned using MTL.

**Fig 3:**
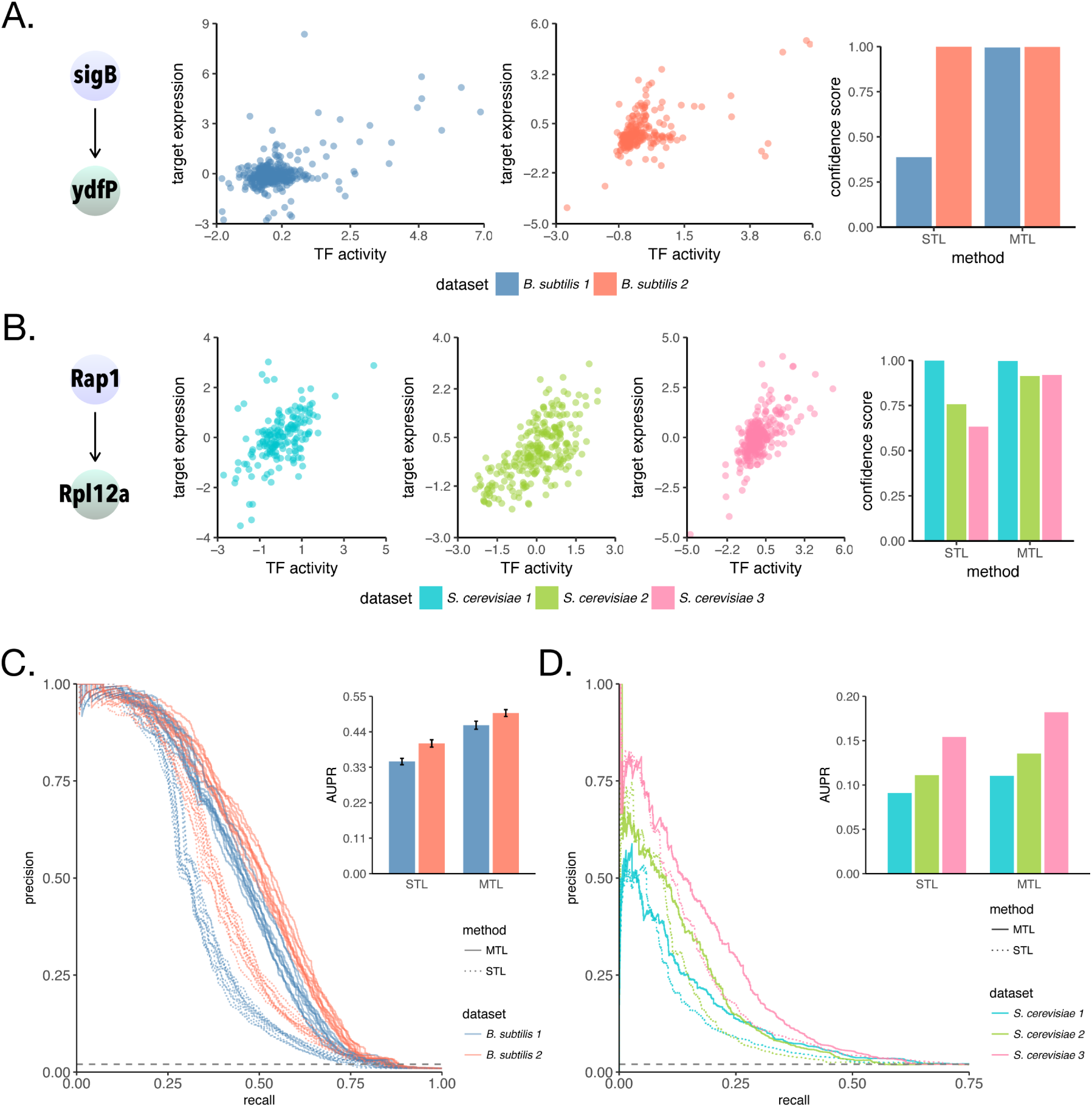
Multitask learning improves accuracy of inferred networks. (A) Relationship between TF activity and target expression in *B. subtilis* 1 (blue) and in B. subtilis 2 (orange), and corresponding STL and MTL inferred confidence scores for an example of an interaction in the *B. subtilis* gold-standard, *sigB* to *ydfP*. (B) as shown in (A), but for an interaction in the *S. cerevisiae* gold-standard, Rap1 to Rpl12a. (C) Precision-recall curves assessing accuracy of network models inferred for individual *B. subtilis* datasets against a leave-out set of interactions. Barplot show mean area under precision-recall curve (AUPR) for each method and dataset. Error bars show the standard deviation across 10 splits of the gold-standard into prior and evaluation set. (D) Precision-recall curves assessing accuracy of network models inferred for individual *S. cerevisiae* networks, with the difference that the prior is from an independent source (no splits or replicates).

In order to evaluate the overall quality of the inferred networks, we use area under precision-recall curves (AUPR) [16], widely used to quantify a classifier’s ability to distinguish two classes and to rank predictions. Networks learned using MTL are significantly more accurate than networks learned using the STL approach. For *B. subtilis* (Fig 3D), we observe around a 30% gain in AUPR for both datasets, indicating significant complementarity between the datasets. For *S. cerevisiae* (Fig 3E), we observe a clear increase in performance for networks inferred for every dataset, indicating that our method is very robust to both data heterogeneity and potential false edges derived from chromatin accessibility in the prior. These experiments were also performed using TF expression as covariates, instead of TF activities, and those results are shown at (Fig S1A, B). Although we recommend using TFA for the organisms here tested, MTL also improves the performance for each dataset-specific network in this scenario.

### Benefits of multitask learning exceed those from batch-correction and ensemble methods

Next, we asked whether the higher performance of the MTL framework could be achieved by other commonly used data integration strategies, such as batch-correction and ensemble methods. Ensemble methods include several algebraic combinations of predictions from separate classifiers trained within a single-domain (sum, mean, maximum, minimum [64]). To address this question, we evaluated networks inferred using all available data. First, we combined regulatory models inferred for each dataset either through STL or MTL by taking the average rank for each interaction, generating two networks hereafter called STL-C and MTL-C [16]. For each organism, we also merged all datasets into one, and applied ComBat for batch-correction [65], because of its perceived higher performance [66]. We then learn network models from these larger batch-corrected datasets, STL-BC. Both for *B. subtilis* (Fig 4A) and *S. cerevisiae* (Fig 4B), the MTL-C networks outperform the STL-C and STL-BC networks, indicating that cross-dataset information sharing during modelling is a better approach to integrate datasets from different domains. Interestingly, for *B. subtilis*, the STL-BC network has a higher performance than the STL-C network, whereas for yeast we observe the opposite. We speculate that the higher overlap between the conditions in the two *B. subtilis* datasets improved performance of the batch-correction algorithm here used. For yeast, on the other hand, conditions were very different across datasets, and although much new information is gained by merging datasets into one, it is likely that incorrect relationships between genes were induced as an artifact, possibly confounding the inference. Of note, these approaches emphasize the commonalities across datasets, whereas the motivation to use MTL frameworks is to increase statistical power, while maintaining separate models for each dataset, hopefully improving interpretability. These experiments were also performed using TF expression as covariates, instead of TF activities, and those results are shown at (Fig S2A, B). In that case, results hold for yeast, but not for *B. subtilis*.

**Fig 4:**
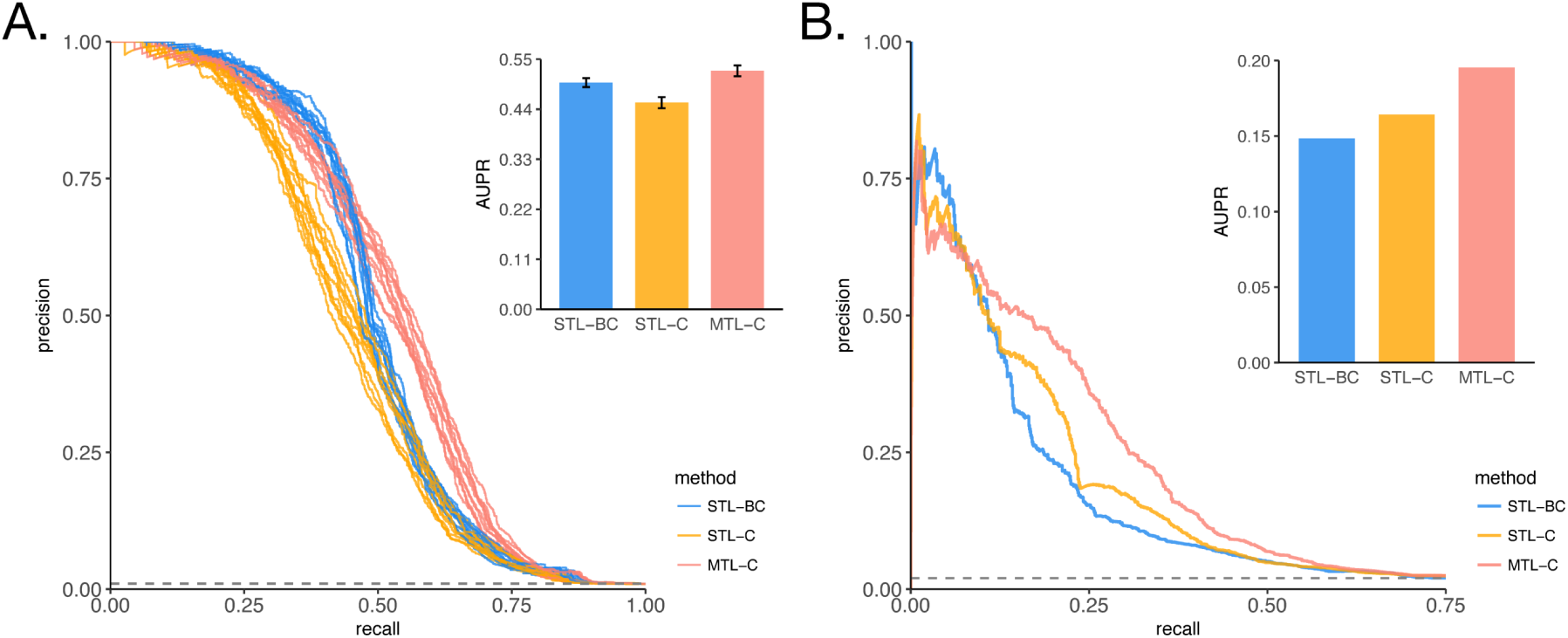
Multitask learning performance boost outweights benefits of other data integration methods. Assessment of accuracy of network models learned using three different data integration strategies, data merging and batch correction (STL-BC), ensemble method combining models learned independently (STL-C), and ensemble method combining models learned jointly (MTL-C). (A) Precision-recall curves for *B. subtilis*, again using a leave-out set of interactions. Barplot show mean area under precision-recall curve (AUPR) for each method. Error bars show the standard deviation across 10 splits of the gold-standard into prior and evaluation set. (B) Precision-recall curves for *S. cerevisiae*, with the difference that the prior is from an independent source (no splits or replicates).

### Our method is robust to increasing prior weights and noise in prior

Because genes are frequently co-regulated, and biological networks are redundant and robust to perturbations, spurious correlations between transcription factors and genes are highly prevalent in gene expression data [67, 68]. To help discriminate true from false interactions, it is essential to incorporate prior information to bias model selection towards interactions with prior knowledge. Indeed, incorporating prior knowledge has been shown to increase accuracy of inferred models in several studies [3, 21, 69].

For example, suppose that two regulators present highly correlated activities, but regulate different sets of genes. A regression-based model would be unable to differentiate between them, and only other sources of information, such as binding evidence nearby a target gene, could help selecting one predictor over the other in a principled way. Thus, we provide an option to integrate prior knowledge to our MTL approach in the model selection step by allowing the user to input a “prior weight”. This weight is used to increase presence of prior interactions to the final model, and should be proportional to the quality of the input prior.

Sources of prior information for the two model organisms used in this study are fundamentally different. The *B. subtilis* prior is high-quality, derived from small-scale experiments, whereas the *S. cerevisiae* prior is noisier, likely with both high false-positive and false-negative rates, derived from high-throughput chromatin accessibility experiments and TF binding motifs. To understand differences in prior influences for the same organism, we also include the yeast gold-standard as a possible source of prior in this analysis. The number of TFs per target gene in the *B. subtilis* (Fig 5A) and the *S. cerevisiae* (Fig 5B) gold-standards (GS) is hardly ever greater than 2, with median of 1, whereas for the chromatin accessibility-derived priors (ATAC) for *S. cerevisiae*, the median is 11 (Fig 5C). A large number of regulators per gene likely indicates a high false-positive rate in the yeast ATAC prior. Given the differences in prior quality, we test the sensitivity of our method to the prior weight parameter. We applied increasing prior weights, and measured how the confidence scores attributed to prior interactions was affected (Fig 5D) for the three source of priors described above. Interestingly, the confidence scores distributions show dependencies on both the prior quality and the prior weights. When the gold-standard interactions for *B. subtilis* and *S. cerevisiae* are used as prior knowledge, they receive significantly higher scores than interactions in the *S. cerevisiae* chromatin accessibility-derived prior, which is proportional to our belief on the quality of the input prior information. Importantly, even when we set the prior weight value to a very high value, such as 10, interactions in the ATAC prior are not pushed to very high confidence scores, suggesting that our method is robust to the presence of false interactions in the prior.

**Fig 5:**
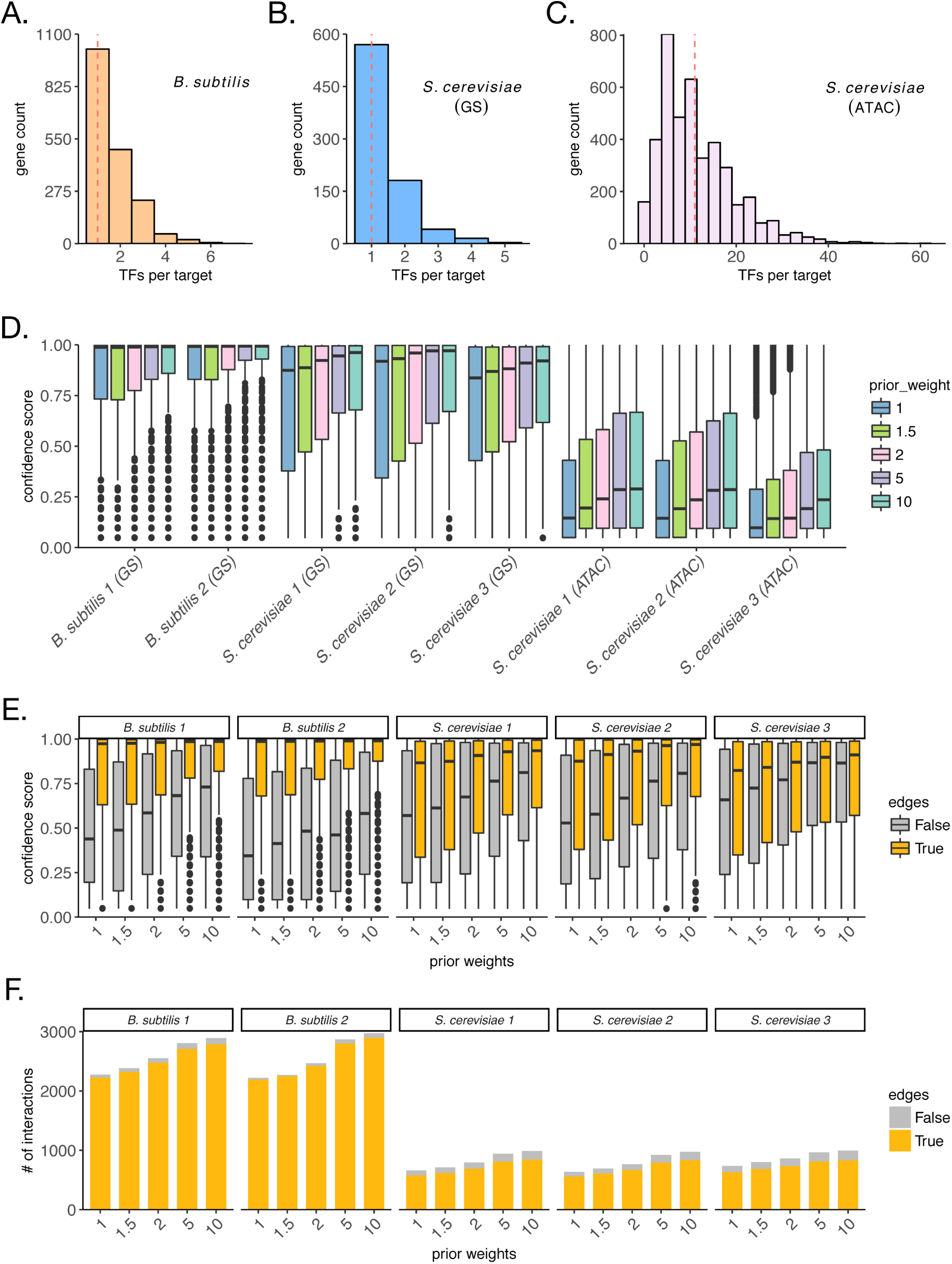
Recovery of prior interactions depends on prior quality and is robust to increasing prior weights. Distribution of number of regulators per target in the *B. subtilis* prior (A), for the *S. cerevisiae* gold-standard (B), and for the *S. cerevisiae* chromatin accessibility-derived priors (C). (D) Distributions of MTL inferred confidence scores for interactions in the prior for each dataset. Different colors show prior weights used, and represent an amount by which interactions in the prior are favored by model selection when compared to interactions without prior information. (E) Distributions of MTL inferred confidence scores for true (yellow) and false (gray) interactions in the prior for each dataset. (F) Counts of MTL inferred interactions with non-zero confidence scores for true (yellow) and false (gray) interactions in the prior for each dataset.

In order to test this hypothesis, we artificially introduced false edges to both the *B. subtilis* and the yeast gold-standards. We added 1 randomly chosen “false” interaction for every 5 true edges in the gold-standard. That affects both TFA estimation and model selection (for prior weights greater than 1). We then ran the inference using the *Inferelator-AMuSR* method with increasing prior weights, and evaluated both the confidence scores of recovered true and false interactions (Fig 5C) as well as the counts of true and false interactions that receive non-zero confidence scores (Fig 5D). For both *B. subtilis* and yeast, we notice that confidence scores distributions show dependency on whether edges are true or false, indicating that the method is not overfitting the prior for the majority of datasets, even when prior weights used are as high as 10 (Fig 5C). We speculate that the greater completeness of the *B. subtilis* gold-standard and of the expression datasets make it easier to differentiate true from false prior interactions when compared to yeast. Besides, inferring networks for prokaryotes is regarded as an easier problem [16]. Importantly, we also show the number of non-zero interactions in each of these distributions (Fig 5D). Taken together, these results show that our method is robust to false interactions in the prior, but requires the user to choose an appropriate prior weight for the specific application. As in previous studies [43], in the presence of a gold-standard, we recommend the user to evaluate performance in leave-out sets of interactions to determine the best prior weight to be used. In the absence of a gold-standard, priors are likely to be of lower confidence, and therefore, smaller prior weights should be used.

### Joint network inference is robust to dataset heterogeneity

Because multitask learning approaches are inclined to return models that are more similar to each other, we sought to understand how heterogeneity among datasets affected the inferred networks. Specifically, we quantified the overlap between the networks learned for each dataset for *B. subtilis* and yeast. That is, the number of edges that are unique or shared across networks inferred for each dataset (Fig 6). In this analysis, we consider valid only predictions within a 0.5 precision cut-off, calculated using only TFs and genes present in the gold-standard. Since the *B. subtilis* datasets share more conditions than the yeast datasets, we hypothesized that the *B. subtilis* networks would have a higher overlap than the yeast networks. As expected, we observe that about 40% of the total edges are shared among two *B. subtilis* networks (Fig 6A), whereas for yeast only about 27% (Fig 6B) and 22% (Fig 6C), using gold-standard and chromatin accessibility-derived priors respectively, of the total number of edges is shared by at least two of the three inferred networks. Therefore, our approach for joint inference is robust to cross-dataset influences, preserving relative uniqueness when datasets are more heterogeneous.

**Fig 6:**
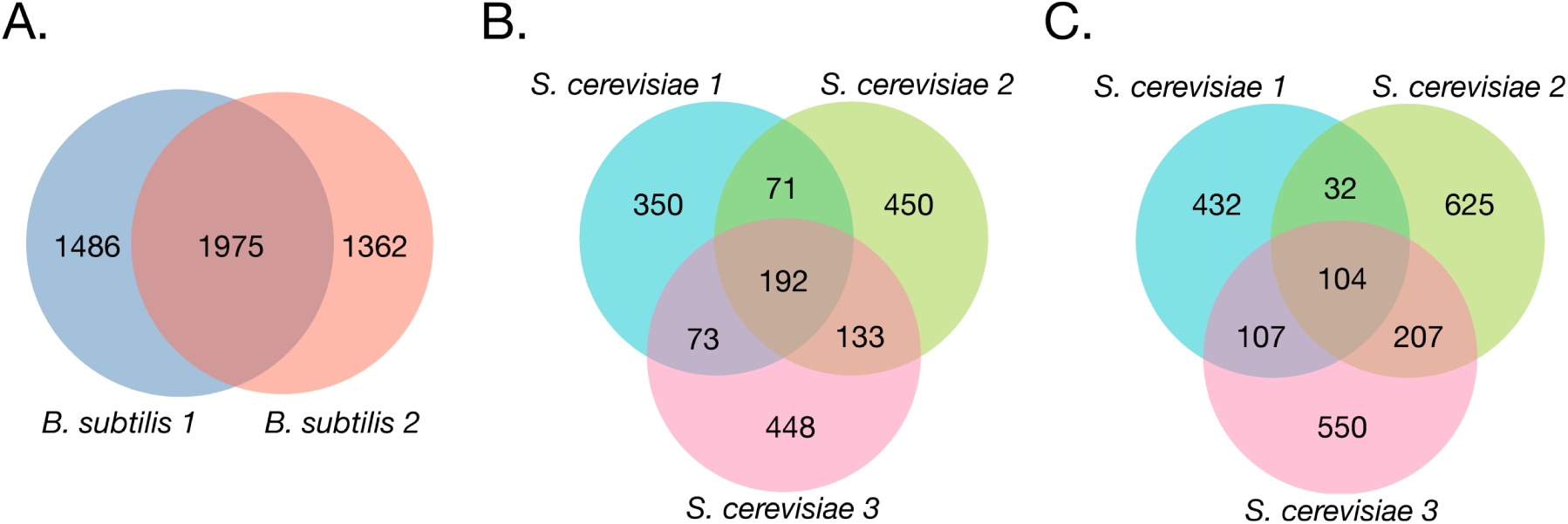
Overlap of edges in inferred networks is higher for *B*. *subtilis* than for *S*. *cerevisiae*. Edges overlap across networks inferred using multitask learning for *B. subtilis* (prior weight of 1.0) (A), for *S. cerevisiae* (using the gold-standard as priors) (B), for *S. cerevisiae* (using the chromatin accessibility-derived priors) (C).

## Discussion

In this study, we presented a multitask learning approach for joint inference of gene regulatory networks across multiple expression datasets that improves performance and biological interpretation by factoring network models derived from multiple datasets into conserved and dataset-specific components. Our approach is designed to leverage cross-dataset commonalities while preserving relevant differences. While other multitask methods for network inference penalize for differences in model coefficients across datasets [25–28, 30], our method leverages shared underlying topology rather than the influence of TFs on targets. We expect this method to be more robust, because, in living cells, a TF’s influence on a gene’s expression can change in different conditions. In addition, previous methods either deal with dataset-specific interactions [25], or apply proper sparsity inducing regularization penalties [26–30]. Our approach, on the other hand, addresses both concerns. Finally, we implemented an additional feature to allow for incorporation of prior knowledge on network topology in the model selection step.

Using two different model organisms, *B. subtilis* and *S. cerevisiae*, we show that joint inference results in accurate network models. We also show that multitask learning leads to more accurate models than other data integration strategies, such as batch-correction and combining fitted models. Generally, the benefits of multitask learning are more obvious when task overlap is high and datasets are slightly under-sampled [34]. Our results support this principle, as the overall performance increase of multitask network inference for *B. subtilis* is more pronounced than for *S. cerevisiae*, which datasets sample more heterogeneous conditions. Therefore, to benefit from this approach, defining input datasets that share underlying regulatory mechanisms is essential and user-defined.

A key question here, that requires future work, is the partitioning of data into separate datasets. Here we use the boundaries afforded by previous study designs: we use data from two platforms and two strains for *B. subtilis* (a fairly natural boundary) and the separation between studies by different groups (again using different technologies) in yeast. We choose these partitions to illustrate robustness to the more common sources of batch effect in meta-analysis. In the future, we expect that multitask methods in this domain will integrate dataset partition estimation (which data go in which bucket) with network inference. Such methods would ideally be able to estimate task similarity, taking into account principles of regulatory biology, and apply a weighted approach to information sharing. In addition, a key avenue for future work will be to adapt this method to multi-species studies. Examples of high biological and biomedical interest include joint inferences across model systems and organisms of primary interest (for example data-sets that include mouse and human data collected for similar cell types in similar conditions). These results (and previous work on many fronts [7, 25, 70]) suggest that this method would perform well in this setting. Nevertheless, because of the increasing practice of data sharing in Biology, we speculate that cross-study inference methods will be largely valuable in the near future, being able to learn more robust and generalizable hypotheses and concepts. Although we present this method as an alternative to batch correction, we should point out that there are many uses to batch correction that fall outside of the scope of network inference, and our results do not lessen the applicability of batch correction methods to these many tasks. There is still great value in properly balancing experimental designs when possible to allow for the estimation of specific gene- and condition-wise batch effects. Experiments where we interact MTL learning with properly balanced designs and quality batch correction are not provided here, but would be superior. Thus, the results here should be strictly interpreted in the context of network inference, pathway inference, and modeling interactions.

## Methods

### Expression data selection, preprocessing and batch-correction

For *B. subtilis*, we downloaded normalized expression datasets from the previously published network study by Arrieta-Ortiz *et al* [17]. *Both datasets are available at GEO, B. subtilis 1* with accession number GSE67023 [17] and *B. subtilis 2* with accession number GSE27219 [52]. For yeast, we downloaded expression datasets from the SPELL database, where hundreds of re-processed gene expression data is available for this organism. In particular, we selected three datasets from separate studies based on the number of samples, within-dataset condition diversity, and cross-dataset condition overlap (such as nutrient-limited stress). *S. cerevisiae 1* and *S. cerevisiae 2* are also available at GEO at accession numbers GSE11452 [54] and GSE9376 [55]. *S. cerevisiae 3* does not have a GEO accession number, and was collected in a custom spotted microarray [56]. For network inference, we only kept genes present in all datasets, resulting in 3780 and 4614 genes for *B. subtilis* and for yeast respectively. In order to join merge, for comparison, we consider each dataset to be a separate batch, since they were generated in different labs as part of separate studies, and applied ComBat for batch-correction using default parameters and no reference to experimental designs [65].

### Building priors from chromatin accessibility

#### ATAC-seq data download, processing, and peak calling

We downloaded chromatin accessibility data for *S. cerevisiae* from the European Nucleotide Archive (PRJNA276699) [71, 72]. Reads were mapped to the sacCer3 genome (iGenomes, UCSC) using bowtie2 [73] with the options –very-sensitive –maxins 2000. Reads with low mapping quality (MAPQ *<* 30), or that mapped to mitochondrial DNA were removed. Duplicates were removed using Picard. Reads mapping the forward strand were offset by +4 bp, and reads mapping to the reverse strand −4 bp. Accessible regions were called using MACS2 [74] with the options –qvalue 0.01 –gsize 12100000 –nomodel –shift 20 –extsize 40. We defined the union of peaks called in any the ATAC-seq samples as the set of putative regulatory regions.

#### Motifs download, assignment to target genes, and prior generation

We obtained a set of expert-curated motifs for *S. cerevisiae* containing position frequency matrices for yeast transcription factors from The Yeast Transcription Factor Specificity Compendium motifs (YeTFaSCo) [75]. Then, we scanned the whole yeast genome for occurrences of motifs using FIMO with p-value cutoff 1e-4 [59], and kept motifs that intersected with putative regulatory regions. Each motif was then assigned to the gene with closest downstream transcription start site. Gene annotations were obtained from the Saccharomyces Genome Database (SGD) [76]. A list of putative regulators was downloaded from the YeastMine database [62], and then generated a targets-by-regulators matrix (prior) where entries are the count of motifs for a particular regulator assigned to each gene. Finally, we multiplied entries for repressors by −1.

### Network inference

We approach network inference by modeling gene expression as a weighted sum of the activities of transcription factors [17, 36]. Our goal is to learn these weights from gene expression data as accurately as possible. In this section, we explain our core model of gene regulation, and of transcription factor activities, and state our assumptions. We also describe how we extend our framework to support learning of multiple networks simultaneously, and integration of prior knowledge on network structure. Finally, we explain how we rank predicted interactions which is used to evaluate the ability of these methods to recover the known underlying network.

#### Core model

We model the expression of a gene *i* at condition *j, X*_*i,j*_, as the weighted sum of the activities of each transcription factor *k* at condition *j, A*_*k,j*_ [17, 43]. Note that although several methods use transcription factor expression as an approximation for its activity, we explicitly estimate these values from expression data and a set of a prior known interactions. Strength and direction (activation or repression) of a regulatory interaction between transcription factor *k* and gene *i* is represented by *i, k*. At steady state, we assume:

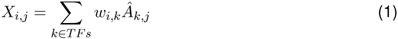

For time-series, we reason that there is a delay *τ* between transcription factor activities and resulting changes in target gene expression [43]. Given expression of a gene *i* in time *t*_*n*_, 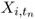, and activity of transcription factor *k* at time *t*_*n-τ*_, 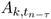, we assume:

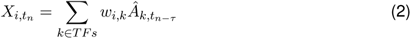

If time *tn - τ* is not available in the expression data, linear interpolation is used to fit 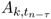.

Finally, because we expect each gene to be regulated by only a few transcription factors, we seek a sparse solution for *w*. That is, a solution in which most entries in *w* are zero. Of note, we set *τ* = 15 for *B. subtilis* [17]. *For S. cerevisiae*, all experiments are considered steady-state.

#### Estimating transcription factor activities (TFA)

We use the expression of known targets of a transcription factor to estimate its activity. From a set of prior interactions, we build a connectivity matrix *P*, where entries represent known activation, *P*_*i,k*_ = 1, or repression, *P*_*i,k*_ = −1, of gene *i* by transcription factor *k*. If no known interaction, *P*_*i,k*_ = 0. We assume that the expression of a gene can be written as a linear combination of the activities of its prior known regulators [17].

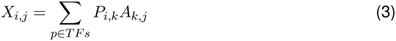

In case of time-series experiments, we use the expression of genes at time *t*_*n*+*τ/*2_, 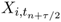, to inform the activities at time *t*_*n*_, *A*_*n*_. Note that for estimating activities, the time delay used is *τ/*2. Again, linear interpolation is used to estimate 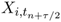 if gene expression at *t*_*n*+*τ/*2_ was not measured experimentally [17].

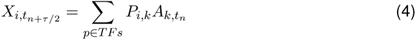

In matrix form, both time-series and steady-state equations can be written as *X* = *PA*. Since there are more target genes than regulators *i > p*, this is an over-determined system, and thus has no solution, so we approximate *A* by finding *Â* that minimizes 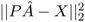. The solution is given by *Â* = *P*^*\**^*X*, where *P*^*\**^ = (*P*^*T*^ *P)*^-1^*P*^*T*^, the pseudo-inverse of *P*. Finally, for transcription factors with no targets in *P*, we use the measured expression values as proxy for the activities.

#### Learning regression parameters

Given gene expression and activity estimates, the next step is to define a set of regulatory hypotheses for the observed changes in gene expression. For each gene, we find a sparse solution for the regression coefficients where nonzero values indicate the transcription factors that better explain the changes observed in gene expression. In this section, we explain how we learn these parameters from a single dataset (single-task learning) and from multiple (multitask learning).

#### Single-task learning using lasso regression (*l*_1_)

The lasso (least absolute selection and shrinkage operator) is a method that performs both shrinkage of the regression coefficients and model selection [49]. That is, it shrinks regression coefficients towards zero, while setting some of them to exactly zero. It does so by adding a penalty on the sum of the absolute values of the estimated regression coefficients. Let *Â* be the activities matrix, *X*_*i*_ the expression values for gene *i*, and *w* the vector of coefficients, lasso estimates are given by:

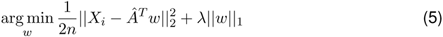

where ‖*w*‖_1_ = ∑_*k*_ |*w*_*k*_|. When minimizing the above function, we seek a good fit while subject to a “budget” on the regression coefficients. The hyper-parameter *λ* controls how much weight to put on the *l*_1_ penalty. The lasso became very popular in the last decade, because it reduces overfitting and automatically performs variable selection. We choose the lasso as a single-task baseline because it is equivalent to the *S* matrix in the multitask case (see below), but with independent choice of sparsity parameter for each dataset.

#### Multitask learning using sparse block-sparse regression (*l*_1_*/l*_1_ **+** *l*_1_*/l*_*∞*_)

We extend our core model to the multiple linear regression setting to enable simultaneous parameter estimation. Here we represent regression parameters for a single gene *i* as a matrix *W*, where rows are transcription factors *k* and columns are networks (or datasets) *d*. We seek to learn the support *Supp*(*W)*, where nonzero entries *W*_*k,d*_ represent a regulatory interaction between transcription factor *k* and gene *i* for network from dataset *d*.

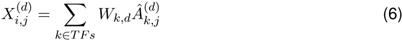

For a given gene *i*, we could assume that the same regulatory network underlies the expression data in all datasets *d*. That is, rows in *W* are either completely non-zero or zero. Since a different set of experiments may have different regulatory patterns, this could be a very strong assumption. A more realistic scenario would be that for each gene *i*, certain regulators are relevant to regulatory models for all datasets *d*, while others may be selected independently by each model *d*. Thus, some rows in the parameter matrix *W* are entirely nonzero or zero, while others do not follow any particular rule. In this scenario, the main challenge is that a single structural constraint such as row-sparsity does not capture the structure of the parameter matrix *W*. For these problems, a solution is to model the parameter matrix as the combination of structurally constrained parameters [77].

As proposed by Jalali et al. [32], we learn the regression coefficients by decomposing *W* into *B* and *S*, that encode similarities and differences between regulatory models respectively. This representation combines a block-regularization penalty on *B* enforcing row-sparsity ‖*B*‖_1,*∞*_ = ∑_*k*_ ‖*B*_*k*_‖_*∞*_, where ‖*B*_*k*_‖_*∞*_:= max_*d*_ |*B*_*k,d*_| (as the one from the previous section), and an elementwise penalty on *S* allowing for deviation across regulatory models for each dataset ‖*S*‖_1,1_ = ∑_*k,d*_ |*S*_*k,d*_|. The goal is to leverage any parameter overlap between models through *B*, while accommodating the differences through *S*. We obtain an estimate for *Ŵ* by solving the following optimization problem:

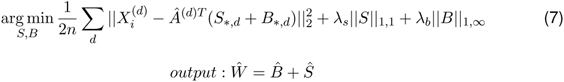

#### Incorporating prior knowledge using the adaptive lasso

We incorporate prior knowledge by differential shrinkage of regression parameters in *S* through the adaptive lasso [33]. We choose to apply this only to the *S* component, because we wanted to allow the user to input different priors for each dataset if so desired. Intuitively, we penalize less interactions present in the prior network. Let Φ be a matrix of regulators *k* by datasets *d*, such that entries Φ_*k,d*_ are inversely proportional to our prior confidence on the interaction between regulator *k* and gene *i* for dataset *d*. We then optimize the following objective:

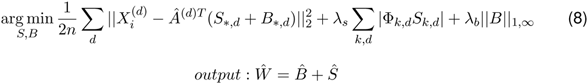

We implement this by scaling *λ*_*s*_ by Φ, then the penalty applied to *S*_*k,d*_ becomes Φ_*k,d*_*λ*_*s*_. In the extreme Φ_*k,d*_ = 0, the regulator *k* is not penalized and will be necessarily included in the final model for dataset *d*. In practice, the algorithm accepts an input prior weight *ρ* ≥ 1 that is used to generate the matrix Φ. We apply the complexity-penalty reduction afforded by Φ_*k,d*_ to *Ŝ* and not 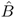 as this choice penalizes unique terms, creating the correct behavior of encouraging model differences that are in accord with orthogonal data as expressed in the network-prior. This choice is also in accord with the interpretation of the prior as valid in one, but not necessarily all, conditions. If regulator *k* is in the prior for dataset *d*, then Φ_*k,d*_ = 1*/ρ*, otherwise Φ_*k,d*_ = 1. Finally, we rescale Φ_*\*,d*_ to sum to the number of predictors *k*. Note that each network model accepts its own set of priors.

#### Model selection

As proposed by Jalali et al. [32], for MTL, we set 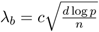, with *n* being the number of samples, *d* being the number of datasets, and search for *c* in the logarithmic interval [0.01, 10]. For each *λ*_*b*_, we look for *λ*_*s*_ that satisfy 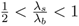. We choose the optimal combination (*λ*_*s*_, *λ*_*b*_) that minimizes the extended Bayesian information criterion (EBIC) [78], here defined as:

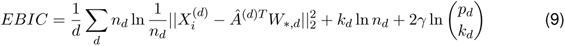

with *k*_*d*_ being the number of nonzero predictors in *W* for model *d*, and 0 *≤ γ ≤* 1. Note that for *γ* = 0, we recover the original BIC. Whereas for *γ >* 0, the EBIC scales with the predictor space *k* making it particularly appropriate for scenarios where *p ≫ n*, often encountered in biological network inference projects. In this study, we set *γ* = 1. For STL, we use the same EBIC measure, but we calculate it for each dataset separately. Importantly, model selection using EBIC is significantly faster than when compared to re-sampling approaches, such as cross-validation or stability selection [79]. Cross-validation, for example, was previously reported as an impediment for multitask learning in large-scale network inference due computational feasibility [29].

#### Implementation

We implemented the MTL objective function using cyclical coordinate descent with covariance updates. That is, at each iteration of the algorithm we cycle through the predictors (coordinates), and minimize the objective at each predictor *k* while keeping the others fixed. Briefly, for a given (*λ*_*s*_, *λ*_*b*_), we update entries in *S* and *B* respectively, while keeping other values in these matrices unchanged, for several iterations until convergence. First, we update values in *S* by:

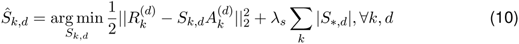

with 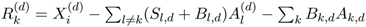, being the partial residual vector. Intuitively, we remove effect of the previous coefficient value for *S*_*k,d*_, while keeping *B*_*k,d*_ unchanged and measure how it changes the residuals. This represents a measure of how important that feature is to the prediction, and contributes to the decision of whether a feature is pushed towards zero or not by the lasso penalty. For *λ*_*s*_ = 0, we can find the least squares update, 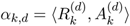, and re-write as

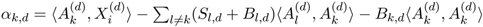. This formulation can be optimized much quicker using the covariance updates explained below.

Then, we update 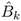, which represents an entire row in *B*, by:

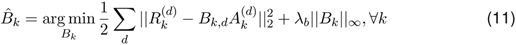

with 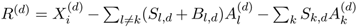, being the partial residual vector for this case. In this case, we keep the value *S*_*k,d*_ unchanged, and set *B*_*k,d*_ to zero. Similarly, we remove effects from previous *B*_*k,d*_ and evaluate how this feature is for the prediction; this then contributes to the decision of whether this entire row is sent to zero by the infinity norm penalty. For *λ*_*b*_ = 0, we can find the least squares update, 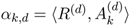, which can be re-written as 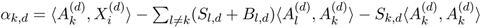. Finally, we apply soft-thresholding to penalize the least-squares updates.

Using these formulations for the updates, we can use the idea of covariance updates [50, 80], where the cross-products *A*^*T*^ *A* and *A*^*T*^ *X* are stored in separate matrices and reused at every iteration. Because these cross-products correspond to over 95% of computation time, this trick decreases runtime significantly. To further decrease runtime, we also employ warm starts when searching for optimal penalty values (*λ*_*s*_, *λ*_*b*_) [80]. Additionally, since we infer regulators for each gene separately, we can parallelize calculations by gene.

### Estimating prediction confidence scores

For each predicted interaction we compute a confidence score that represents how well a predictor explains the expression data, and a measure of prediction stability. As previously proposed [17, 43], we calculate confidence scores for each interaction by:

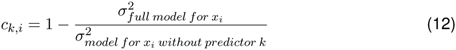

where *σ*^2^ equals the variance of the residuals for the models, with and without predictor *k*. The score *c*_*k,i*_ is proportional to how much removing regulator *k* from gene *i* set of predictors decreases model fit. To measure stability, we perform the inference across multiple bootstraps of the expression data (we used 20 bootstraps for both *B. subtilis* and yeast), rank-average the interactions across all bootstraps [16, 43], and re-scale the ranking between 0 and 1 to output a final ranked list of regulatory hypotheses.

## Implementation and Availability

The *Inferelator-AMuSR* code and example datasets are available at https://github.com/simonsfoundation/multitask_inferelator/tree/AMuSR.

## Supplemental Figure Legends

**Fig S1:**
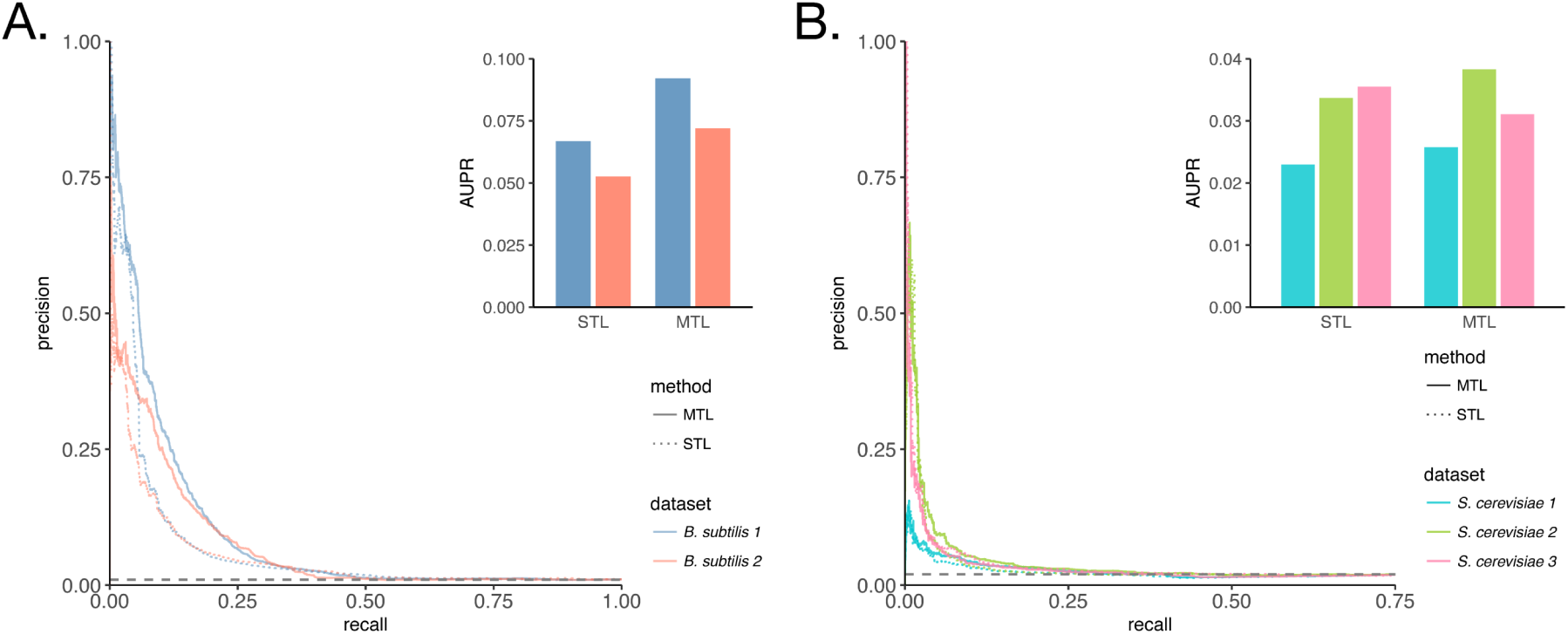
Multitask learning (without TF activities) improves accuracy of inferred networks. (A) Precision-recall curves assessing accuracy of network models inferred without TF activities for individual *B. subtilis* datasets against the whole gold-standard set of interactions. Networks Barplot show mean area under precision-recall curve (AUPR) for each method and dataset. (B) Precision-recall curves assessing accuracy of network models inferred without TF activities for individual *S. cerevisiae* networks, with the difference that priors are derived from chromatin accessibility data.

**Fig S2:**
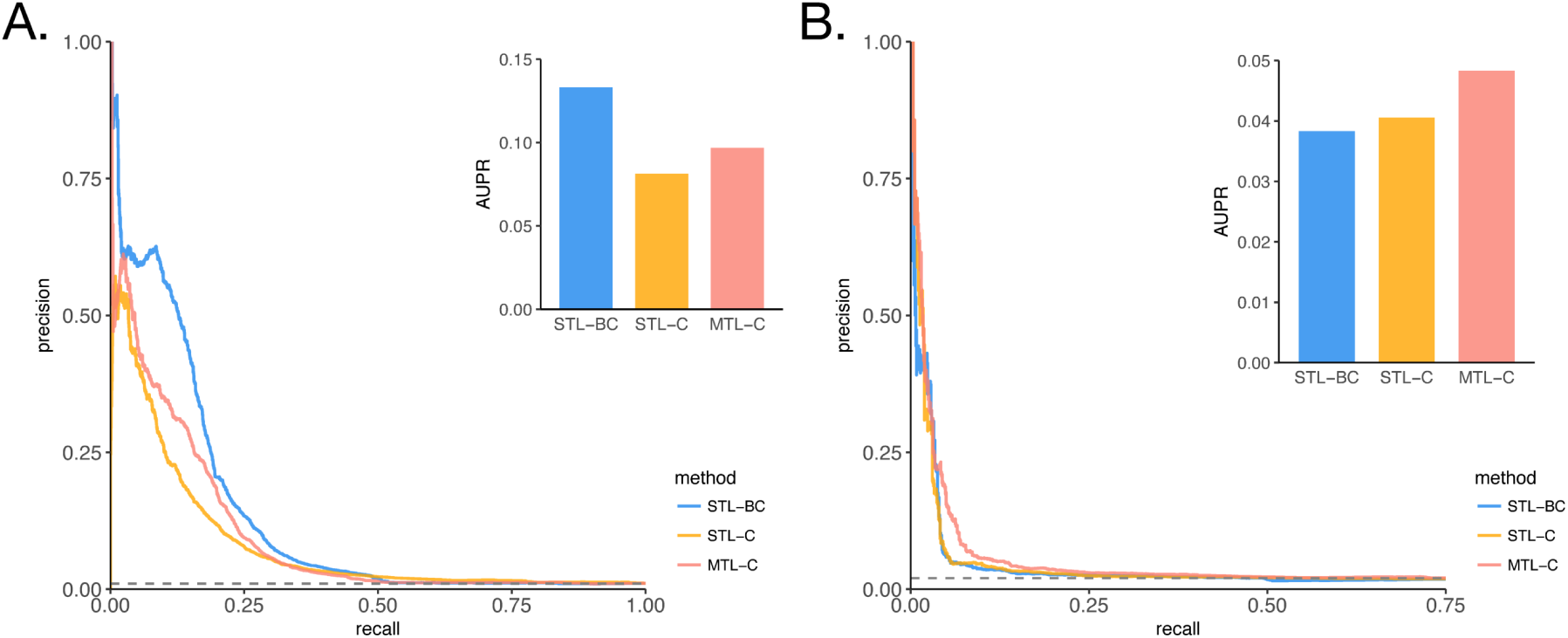
Multitask learning (without TF activities) performance boost outweights benefits of other data integration methods for yeast, but not for *B*. *subtilis*. Assessment of accuracy of network models learned using three different data integration strategies, data merging and batch correction (STL-BC), ensemble method combining models learned independently (STL-C), and ensemble method combining models learned jointly (MTL-C). TF expression was used as predictors of gene expression. (A) Precision-recall curves for *B. subtilis*, again using the whole gold-standard set of interactions. Barplot show mean area under precision-recall curve (AUPR) for each method. (B) Precision-recall curves for *S. cerevisiae*, with the difference that priors are derived from chromatin accessibility data.

